# Benchmarking accuracy and precision of intensity-based absolute quantification of protein abundances in *Saccharomyces cerevisiae*

**DOI:** 10.1101/2020.03.23.998237

**Authors:** Benjamín J. Sánchez, Petri-Jaan Lahtvee, Kate Campbell, Sergo Kasvandik, Rosemary Yu, Iván Domenzain, Aleksej Zelezniak, Jens Nielsen

## Abstract

Protein quantification via label-free mass spectrometry (MS) has become an increasingly popular method for determining genome-wide absolute protein abundances. A known caveat of this approach is the poor technical reproducibility, i.e. how consistent the estimations are when the same sample is measured repeatedly. Here, we measured proteomics data for *Saccharomyces cerevisiae* with both biological and inter-batch technical triplicates, to analyze both accuracy and precision of protein quantification via MS. Moreover, we analyzed how these metrics vary when applying different methods for converting MS intensities to absolute protein abundances. We found that a simple normalization and rescaling approach performs as accurately yet more precisely than methods that rely on external standards. Additionally, we show that inter-batch reproducibility is worse than biological reproducibility for all evaluated methods. These results subsequently serve as a benchmark for assessing MS data quality for protein quantification, whilst also underscoring current limitations in this approach.

## Introduction

Mass spectrometry (MS) is currently the main technology for determining genome wide protein copy number per cell, thanks to its high sensitivity, specificity, and multiplexing capacity [1]. Among the different available MS technologies, quantitative label-free methods have become increasingly popular, due to their relative ease of use and cost-effectiveness, particularly when they are compared to more expensive and laborious methods, as using isotope-labeled peptides [2]. In quantitative label-free approaches, normalization of the raw data is a critical step in determining protein absolute estimates [3–6]. Two fundamental metrics for assessing the quality of protein estimates are: (i) accuracy, i.e. how far away from the true value the prediction is, and (ii) precision, i.e. how variable different estimates are when the same measurement is repeated (also referred to as reproducibility).

When estimating absolute (non-relative) protein abundances using MS, there are several factors that affect the precision and accuracy of quantification: i) the intrinsic biological nature of the proteome, which spans the dynamic range of several orders of magnitude in protein abundances; ii) the physicochemical nature of amino acids, as peptide molecules have different ionization properties (i.e. two similarly abundant molecules might have different responses); and iii) the differences in MS instrumentation (e.g. Orbitraps versus time-of-flight instruments), chromatography and experimental protocols. All of the above factors yield only modest results in MS-based analyses when comparing predictions to the true protein concentrations values [7–9], and a large level of variability across different studies [7,10,11].

Studies that compute absolute protein abundance commonly address biological reproducibility by running biological replicates in the same MS batch [7,12,13]. Awareness however of the impact of the MS instrument, i.e. technical reproducibility, has been less studied. This can be determined by running the same biological sample in the same batch [14], or in separate batches [15]; the latter often referred to as “the batch effect”. As different normalization/scaling methods can be used to estimate protein abundance from raw MS intensities [16], it is interesting to study how these methods propagate the inter-batch technical variability into uncertainty in the final protein abundance estimates. In this study, we analyze both accuracy and technical precision of intensity-based absolute quantification of a proteomics dataset from *S. cerevisiae*, and show how prediction quality can be improved using different normalization/scaling methods. In particular, we show that a simple rescaling method [5] performs as accurately but more precisely than alternatives that rely on the use of costly external standards.

## Methods

We generated a proteomics dataset using the *S. cerevisiae*’s strain CEN.PK113-7D, with multiple biological and technical replicates. Samples were obtained from aerobic glucose-limited chemostats at a dilution rate of 0.1 h^-1^, in triplicate, and mixed with an internal standard (IS), using stable isotope labeling by amino acids in cell culture (SILAC). Here, a lysine auxotrophic strain was grown in medium supplemented with double labelled heavy ^15^N, ^13^C-lysine (Cambridge Isotope Laboratories Inc.); samples were then mixed in a 1:1 ratio with each of the other non-labelled (“light”) samples. The IS was also mixed with an external standard of known concentrations, in a ratio of 6:1.1. The external standard used here was the Proteomics Dynamic Range Standard Set (UPS2) mix (Merck), consisting of 48 human proteins in a dynamic concentration range from 500 amoles to 50 pmoles. All mixed samples were stored at −80°C until analysis, wherein they were similarly processed, to isolate variability from the biology and the MS equipment, and not from other sources such as sample preparation differences.

For proteome identification, samples were digested with 1:50 LysC overnight at room temperature. Peptides were then separated on an Ultimate 3000 RSLCnano system (Dionex), eluted to a Q Exactive Plus (Thermo Fisher Scientific) tandem mass spectrometer, and identified with the MaxQuant 1.4.0.8 software package [17], maintaining the peptide-spectrum match and the protein false discovery rate below 1% using a target-decoy approach. Each sample was measured six times: on three separate batches of the MS instrument (with a time difference of 12 and 30 days), and each time twice, using Top5 and Top10 data-dependent acquisition strategies, wherein only the top five or ten highest intensity peptide peaks per one MS full scan were selected for MS/MS analysis, respectively.

Using the described data as a reference, we then evaluated the ability of four different methods for transforming the MS intensity computed by MaxQuant (which corresponds to the sum of all associated peptide intensities) to protein abundances of the internal standard. The first method, known as intensity based absolute quantification (iBAQ) [3], normalizes each protein MS intensity by the corresponding number of theoretically observable peptides, then infers the abundances of each internal standard protein using a linear model generated from the external standard (normalized protein MS intensity vs. known protein quantities). As this method yields abundances that do not always add up to equal amounts of protein injected per sample (Figures S1 – S2), a second method was also assessed that rescales all abundances from iBAQ to equal the total injected mass. The third method tested was the total protein approach (TPA) [18], which bypasses the need for an external standard and instead assumes that the sum of MS intensities of all detected proteins multiplied by the corresponding molecular weights should be proportional to the total amount of protein injected. Finally, the fourth method analyzed is a variation of the TPA method [5], which first normalizes protein intensities with the number of theoretically observable peptides. 

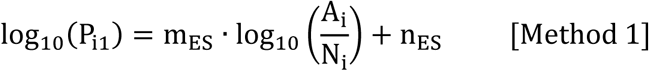

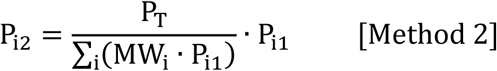

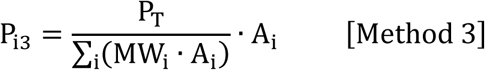

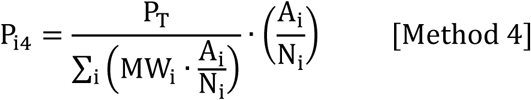

where P_ij_ is the estimated absolute abundance of protein i by method j [fmol/sample], m_ES_ and n_ES_ are the parameters of the external standard curve, A_i_ is the sum of all peptide intensities associated to protein i, N_i_ is the number of theoretically observable peptides for protein i, P_T_ is the total injected protein mass [pg], and MW_i_ is the molecular weight of protein i [kDa]. Finally, for all methods, the sample abundances were calculated based on the corresponding internal (heavy) standard abundance and the normalized H/L ratios obtained from each sample run [19].

## Results and Discussion

Using the generated dataset, we evaluated accuracy and precision of abundances estimated by the four different methods. To evaluate accuracy, we computed the differences as fold changes between the estimated abundances of the external standard proteins detected by the MS (n=31/48) and the known values in the UPS2 mix. Here, Methods 1, 2 and 4 performed similarly, whereas Method 3 had a significantly higher error (Figure 1A, Figure S3). Specifically, more than 50% of protein abundance as estimated by Method 3 deviated from the true value by >2-fold. We further evaluated the accuracy of each Method by testing protein estimates in the ribosome, a protein complex with subunit abundance in equal stoichiometry [20]. Of these subunits, 42 out of 79 were detected (accounting for paralogs) and compared to their median abundance value, as the same abundance for each subunit should be predicted [21]. Once again, Methods 1, 2 and 4 performed similarly, and outperformed Method 3 (Figure 1B, Figures S4 – S5).

**Figure 1:**
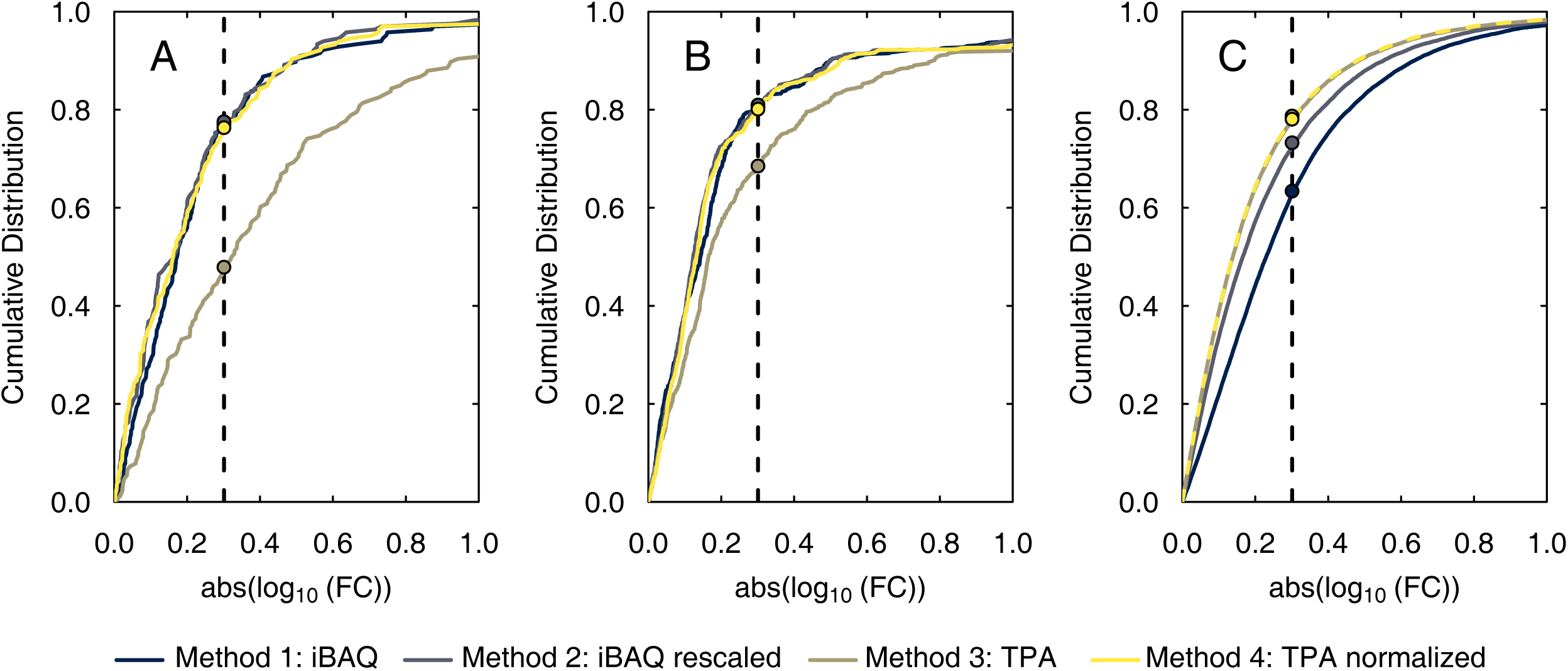
Cumulative distributions of fold changes (FC) between predictions of all four methods, with respect to (A) accuracy (test 1): estimated vs known values of the external standard (N = 167), (B) accuracy (test 2): estimated values vs median estimated value for all ribosomal proteins in the samples (N = 731), and (C) precision: all possible combinations between batches (N = 21,320). A fold change of 2 is indicated with a vertical dashed line.

We then proceeded to evaluate precision, by comparing the protein estimates for each biological sample between all three batches (Figures S6 – S7). A cumulative distribution of all possible fold changes (Figure 1C) showed that Methods 3 and 4 significantly outperformed Methods 1 and 2 (all P-values < 0.001). In particular, TPA-estimated protein abundance varied by less than 2-fold for nearly 80% of all proteins, whereas in the case of iBAQ this was closer to 60%. Higher inter-batch variability of Methods 1 and 2 was observed both for lowly and highly abundant proteins (Figures S8 – S9), and can be explained by the additional variability introduced by the external standard (Figures S10 – S11), that Methods 3 and 4 did not use.

Taking the results of accuracy and precision together (Figure 1), we conclude that the best-performing method is Method 4, which omits use of the external standard and instead rescales the normalized MS intensities to equal the injected sample mass. Even though Methods 1 and 2 perform similarly to Method 4 in terms of accuracy, they are not as precise; and Method 3 although precise, is not as accurate. Therefore, considering that iBAQ involves significant additional cost (purchase of the external standard and additional MS running time), but does not yield better performance, we propose that rescaling normalized MS intensities can be used instead. This method can also be used as a benchmark for assessing the predictive power of alternative approaches for computing absolute protein abundances from MS methods.

It is noteworthy that, despite Method 4 outperforming the other approaches presented here, it is by no means perfect, with ∼20% of estimates still having high technical inter-batch variability, indicative of a batch effect that is not fully resolved. In particular, the variability between biological replicates in the same MS batch (Figure 2A) is considerably lower than the variability between batches of the same biological sample (Figure 2B). This observation can be confirmed with a principal component analysis (Figure 2C), wherein samples cluster based on batches, not biological replicates. Although technical variability becomes lowest when using Method 4 (Figure S6, Table S1), many protein abundances still show to vary in abundance by more than one order of magnitude between batches. This is due to the presence of stochastic and non-linear effects in shotgun proteomics [22,23]; therefore, researchers working with computational methods that rely on absolute protein abundances [24] should be aware of these limitations and interpret results accordingly.

**Figure 2:**
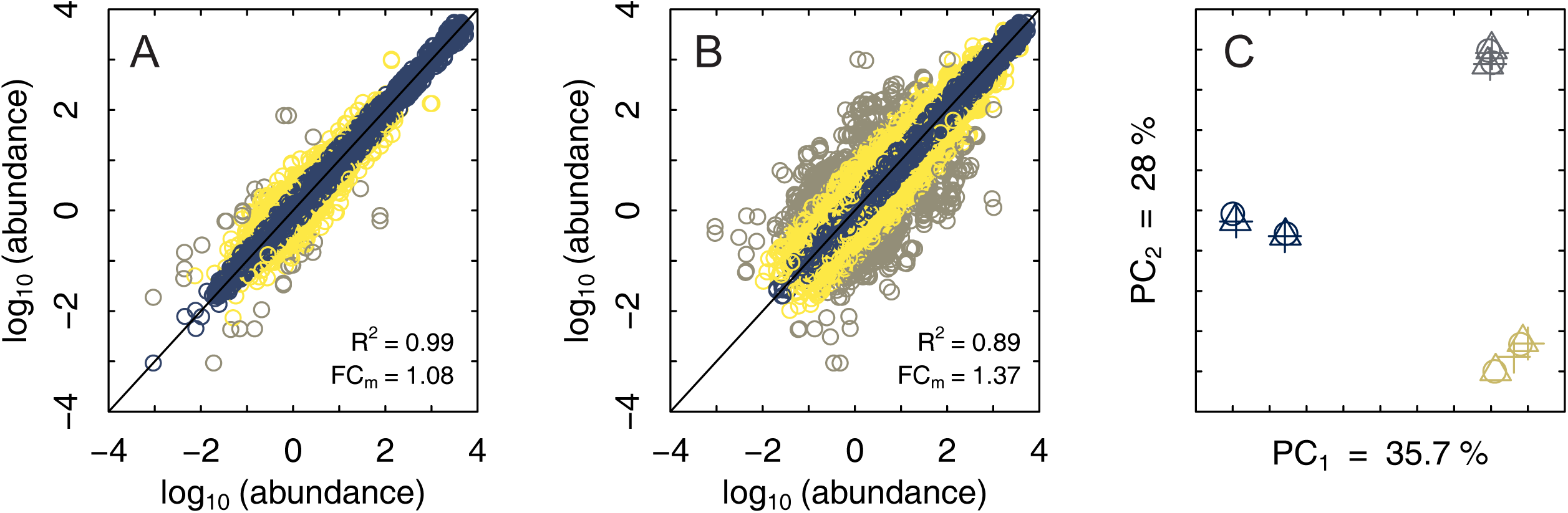
(A-B) Variability between biological replicates (A) and MS batches (B). Fold changes within a 2-fold are shown in blue, between a 2-fold and 10-fold in yellow, and above a 10-fold in gray. The coefficient of determination (R^2^) and the median absolute fold change (FC_m_) are also displayed. (C) Principal component analysis of all samples. Different colors refer to different batches, and different shapes refer to different biological replicates. The amount of variability each of the first 2 components explains is shown as a percentage.

## Conclusion

In conclusion, we present a comprehensive proteomics dataset of yeast, designed for assessment of absolute protein quantification in different biological replicates and batches of samples. Furthermore, we show that a simple method of normalization and rescaling can yield superior results over more complicated and expensive methods such as iBAQ. Finally, as protein intensity is used as input, this method can be used on both pre-existing and future datasets regardless of how intensity values were generated, including labeled or unlabeled methods. We therefore expect both the dataset and method to be useful when assessing accuracy and precision of MS-based proteomics approaches.

All MS data used in this study have been deposited to the ProteomeXchange Consortium via the PRIDE [25] partner repository with the dataset identifier PXD011725. Output tables from MaxQuant, together with all necessary scripts to reproduce the results presented in this study are available at https://github.com/SysBioChalmers/reproduce and have been archived in Zenodo [26].

## Acknowledgments

We would like to thank Dr. Christine Räisänen and Gang Li for reviewing the manuscript. This project has received funding from the European Union’s Horizon 2020 research and innovation program under grant agreement No 686070 and No 668997, the Novo Nordisk Foundation and the Knut and Alice Wallenberg Foundation. BJS and PJL acknowledge financial support from CONICYT (grant #6222/2014) and the Estonian Research Council (grant PUT1488P), respectively.

## Notes

https://github.com/SysBioChalmers/reproduce

## References

[1] Mann, M., Kulak, N.A., Nagaraj, N., Cox, J., Mol. Cell 2013, 49, 583–590.

[2] Calderón-Celis, F., Encinar, J.R., Sanz-Medel, A., Mass Spectrom. Rev. 2017, 37, 715–737.

[3] Schwanhäusser, B., Busse, D., Li, N., Dittmar, G., et al., Nature 2011, 473, 337–342.

[4] Ning, K., Fermin, D., Nesvizhskii, A.I., J. Proteome Res. 2012, 11, 2261–2271.

[5] Wiśniewski, J.R., Ostasiewicz, P., Duś, K., Zielińska, D.F., et al., Mol. Syst. Biol. 2012, 8.

[6] Wiśniewski, J.R., Hein, M.Y., Cox, J., Mann, M., Mol. Cell. Proteomics 2014, 13, 3497–3506.

[7] Arike, L., Valgepea, K., Peil, L., Nahku, R., et al., J. Proteomics 2012, 75, 5437–5448.

[8] Weisser, H., Nahnsen, S., Grossmann, J., Nilse, L., et al., J. Proteome Res. 2013, 12, 1628–1644.

[9] Shuford, C.M., Walters, J.J., Holland, P.M., Sreenivasan, U., et al., Anal. Chem. 2017, 89, 7406–7415.

[10] Lawless, C., Holman, S.W., Brownridge, P., Lanthaler, K., et al., Mol. Cell. Proteomics 2016, 15, 1309–1322.

[11] Ho, B., Baryshnikova, A., Brown, G.W., Cell Syst. 2018, 6, 192-205.e3.

[12] Hukelmann, J.L., Anderson, K.E., Sinclair, L. V, Grzes, K.M., et al., Nat. Immunol. 2016, 17, 104–112.

[13] Lahtvee, P.J., Sánchez, B.J., Smialowska, A., Kasvandik, S., et al., Cell Syst. 2017, 4, 495-504.e5.

[14] Shalit, T., Elinger, D., Savidor, A., Gabashvili, A., Levin, Y., J. Proteome Res. 2015, 14, 1979–1986.

[15] Vowinckel, J., Zelezniak, A., Bruderer, R., Mülleder, M., et al., Sci. Rep. 2018, 8, 1–10.

[16] Rosenberger, G., Ludwig, C., Röst, H.L., Aebersold, R., Malmström, L., Bioinformatics 2014, 30, 2511–2513.

[17] Cox, J., Mann, M., Nat. Biotechnol. 2008, 26, 1367–1372.

[18] Wiśniewski, J.R., Rakus, D., J. Proteomics 2014, 109, 322–331.

[19] Geiger, T., Cox, J., Ostasiewicz, P., Wisniewski, J.R., Mann, M., Nat. Methods 2010, 7, 383–385.

[20] Jenner, L., Melnikov, S., de Loubresse, N.G., Ben-Shem, A., et al., Curr. Opin. Struct. Biol. 2012, 22, 759–767.

[21] Fabre, B., Lambour, T., Bouyssié, D., Menneteau, T., et al., EuPA Open Proteomics 2014, 4, 82–86.

[22] Leek, J.T., Johnson, W.E., Parker, H.S., Jaffe, A.E., Storey, J.D., Bioinformatics 2012, 28, 882–883.

[23] Čuklina, J., Computational Challenges in Biomarker Discovery from High-Throughput Proteomic Data. ETH Zurich, 2018.

[24] Sánchez, B.J., Zhang, C., Nilsson, A., Lahtvee, P., et al., Mol. Syst. Biol. 2017, 13, 935.

[25] Vizcaíno, J.A., Csordas, A., Del-Toro, N., Dianes, J.A., et al., Nucleic Acids Res. 2016, 44, D447–D456.

[26] Sánchez, B.J., 2020, https://doi.org/10.5281/zenodo.3713296.

