## Supplementary Material for "Benchmarking accuracy and precision of intensity-based absolute quantification of protein abundances in *Saccharomyces cerevisiae*"

**References** **14**

### 1 Summary

Here we will go through the typical way of deducing protein abundances [fmol/sample] from SILAC/iBAQ data, and compare it to rescaling values to a fix total protein abundance based on MS intensities, to assess the usefulness of the external standard curve and iBAQ data. The main observation that comes from this is that as MS measurements are so variable, it's impossible to find a unique ES curve, hence normalizing to a fixed total protein abundance is as good as using the "optimal" fit from the ES curve. We can then use the MS intensity directly and bypass both the ES curve and the iBAQ values.

### 2 Loading and pre-processing data

#### 2.1 UPS2

The commercial kit brings 10.6 ug of protein, however only 1.1 ug is injected in the MS.

#### 2.2 iBAQ data

In the iBAQ data (IS+ES) there are 6 samples: measured in 3 batches (`batch1`, `batch2` & `batch3`) and each of them processed with 2 different MS methods `top5` & `top10`. We will have then 6 different ES curves:

- top5_batch1
- top5_batch2
- top5_batch3
- top10_batch1
- top10_batch2
- top10_batch3

In each sample of the iBAQ data there are:

- 6 ug of IS: yeast samples, all marked; i.e. will appear in the heavy fraction (H)
- 1.1 ug of ES: universal protein standard (UPS2) unmarked; i.e. will appear in the light fraction (L)

#### 2.3 SILAC data

There are 18 different samples:

- 3 biological replicates (`R1`, `R2` & `R3`)
- each measured on 3 different batches (`batch1`, `batch2` & `batch3`)
- each estimated with a different MS method (`top5` & `top10`)

Each injected sample consisted of:

- 6 ug of IS (detected in the H fraction)
- 6 ug of actual sample (detected in the L fraction)

#### 2.4 Number of theoretical peptides

We can obtain the number of theoretical peptides for each of the proteins if we remove the UPS2 sequences and label from the MaxQuant search, which leads the software to report only the iBAQ intensities of the proteins. We then divide the total raw intensity with the total iBAQ intensity to get the desired number, and merge this information with the IS and ES data.

#### 2.5 Ribosomal proteins

We used a list of ribosomal genes based on previous work (Jenner et al. 2012).

### 3 Methods evaluated

#### 3.1 Method 1: iBAQ

Method 1 uses the computed iBAQ abundances available in the MaxQuant (Cox and Mann 2008) output file, which are inferred using an ES curve of the UPS2 proteins (in the L fraction) (Schwanhäusser et al. 2011). As the data already comes in fmol/sample, the only thing missing is to use the values from the IS (H fraction) together with the normalized L/H ratios in the SILAC data for getting absolute abundances in each sample of the SILAC data (fmol/sample), by doing:

`abundance(sample) = (L/H)ratio * abundance(IS)`

#### 3.2 Method 2: Rescaling iBAQ values

As iBAQ values don't add up always to the same totals (Figure S1), we should asses the benefits of rescaling all of these values to add up to the injected amounts:

`abundance = (iBAQ abundance)*(injected amount)/(sum of all iBAQ abundances*MW values)`

We now use the absolute abundances from the IS (fmol/sample) to infer absolute abundances in each sample of the SILAC data (fmol/sample).

#### 3.3 Method 3: TPA

The alternative: To skip iBAQ values and ES curves entirely, and to assume all MS intensities summed up together (mass-wise) are proportional to the injected amount in ug. This is known as the total protein approach (TPA) (Wiśniewski and Rakus 2014):

`abundance = (MS intensity)*(injected amount)/(sum of all MS intensities*MW values)`

#### 3.4 Method 4: Normalized TPA

We will also try out to first normalize all MS intensitiy values by the corresponding number of theoretical peptides, to later rescale the data as before (Wiśniewski et al. 2012):

`abundance = (normalized MS intensity)*(injected amount)/(sum of all normalized MS intensities*MW values)`

Note that for methods 3 and 4 we have essentially created a linear model:

`abundance = m*intensity`, where `m = (6 ug)/(sum of intensities*MWs)`

`log(abundance) = log(intensity) + log(m)` -> linear model with a = 1 and b = log(m) in the log space.

### 4 Method comparison

#### 4.1 Protein totals

First, let's take a look at the total detected protein amount in each of the 6 samples of IS, for all methods:

```
## [1] "Method 1: iBAQ - 3497 proteins detected"

## [1] "Method 2: iBAQ rescaled - 3497 proteins detected"

## [1] "Method 3: TPA - 3534 proteins detected"

## [1] "Method 4: TPA normalized - 2923 proteins detected"
```

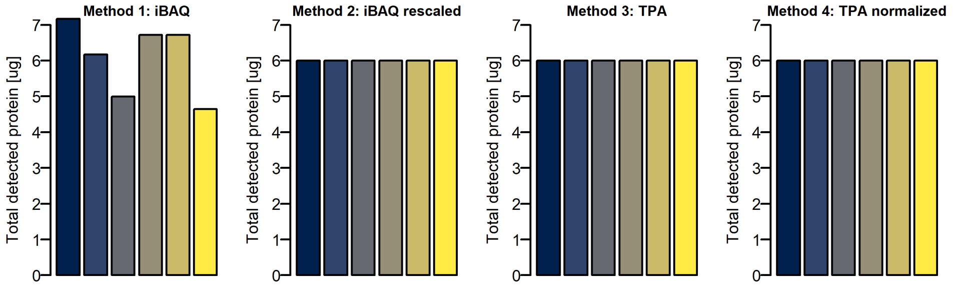

Figure S1: Total detected protein amounts in all 6 IS, according to all methods.

Let's also take a look at the total detected protein amount of each of the 18 samples, colored by the original ES curve used for the calibration (Figure S1):

```
## [1] "Method 1: iBAQ - 1587 proteins detected"

## [1] "Method 2: iBAQ rescaled - 1587 proteins detected"

## [1] "Method 3: TPA - 1587 proteins detected"

## [1] "Method 4: TPA normalized - 1462 proteins detected"
```

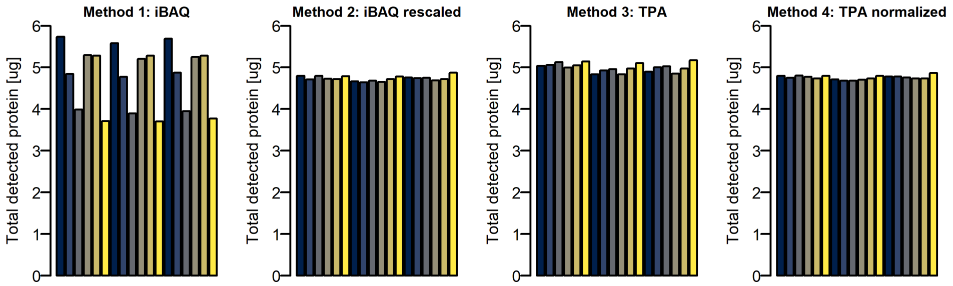

Figure S2: Total detected protein amounts in all 18 samples, according to all methods.

In both figures S1 and S2, the total detected protein seems to vary quite a bit among samples calculated with iBAQ, and as expected this reduces if we rescale (methods 2, 3 and 4). Note that for all cases the total amount of protein detected in the samples is lower than the amount detected in the internal standard, due to more proteins detected in the internal standard (as the latter is a mix of different conditions and not just one sample).

Note as well that the coverage of method 4 is lower than methods 1, 2 and 3. This appears as a limitation of method 4, but it is actually a limitation of the MaxQuant software, which does not provide as an output the number of theoretical peptides for each protein, which led us to have to infer them as described in section 2.4. In fact methods 1 and 2 also employ the number of theoretical peptides, so for any other software this would not be a limitation. In any case, the coverage decrease was very low for the proteins later analyzed: When analyzing accuracy (section 4.2) we only lost one value when comparing UPS2 values (167 → 166) and no values when comparing ribosome stoichiometry (731 → 731). When analyzing precision (section 4.3.1), we only lost 1.2% of values (21,320 → 21,061).

#### 4.2 Evaluating accuracy

Let's now compare accuracy. First, we compare the ES values predicted by each method to the actual UPS2 values:

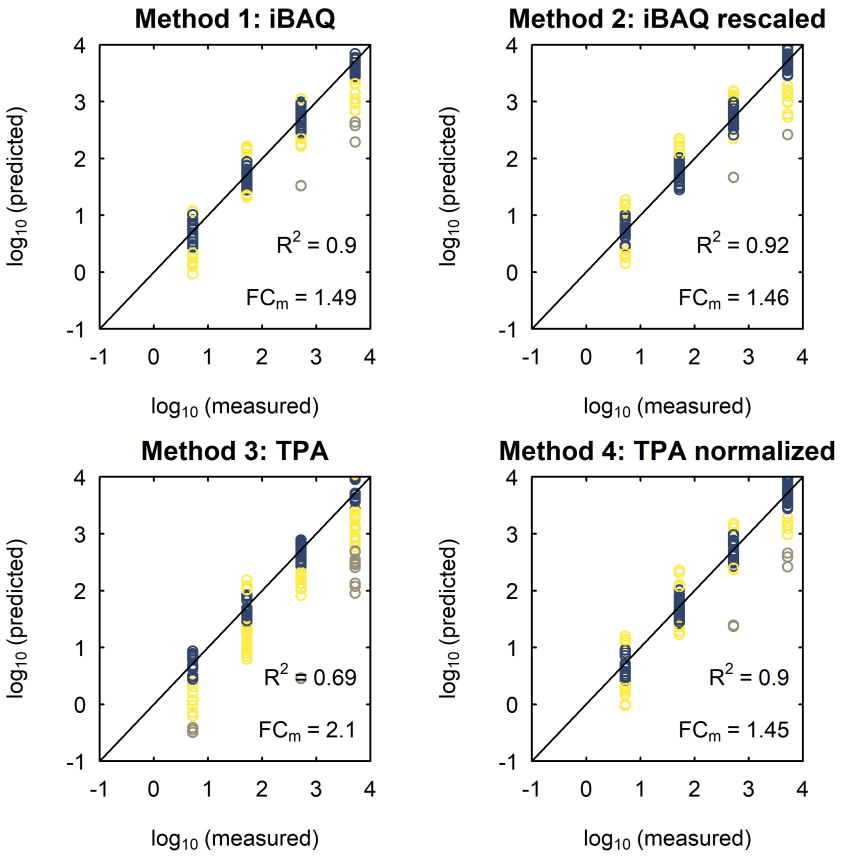

Figure S3: Comparison of predicted Vs real abundance values [fmol/sample] from UPS2, according to all methods. Blue is a FC lower than 2, yellow between 2 and 10, and gray over 10.

We see that all predictions from methods 1, 2 and 4 are very similar; by using ES curves (methods 1-2) we don't gain much prediction power than if we just rescale the normalized data (method 4). However, method 3 performs significantly worse.

Now, let's see how are the predictions of ribosomal subunits stoichiometry. For that first we need to create dataframes with only ribosomal proteins, and then plot for each method the corresponding data:

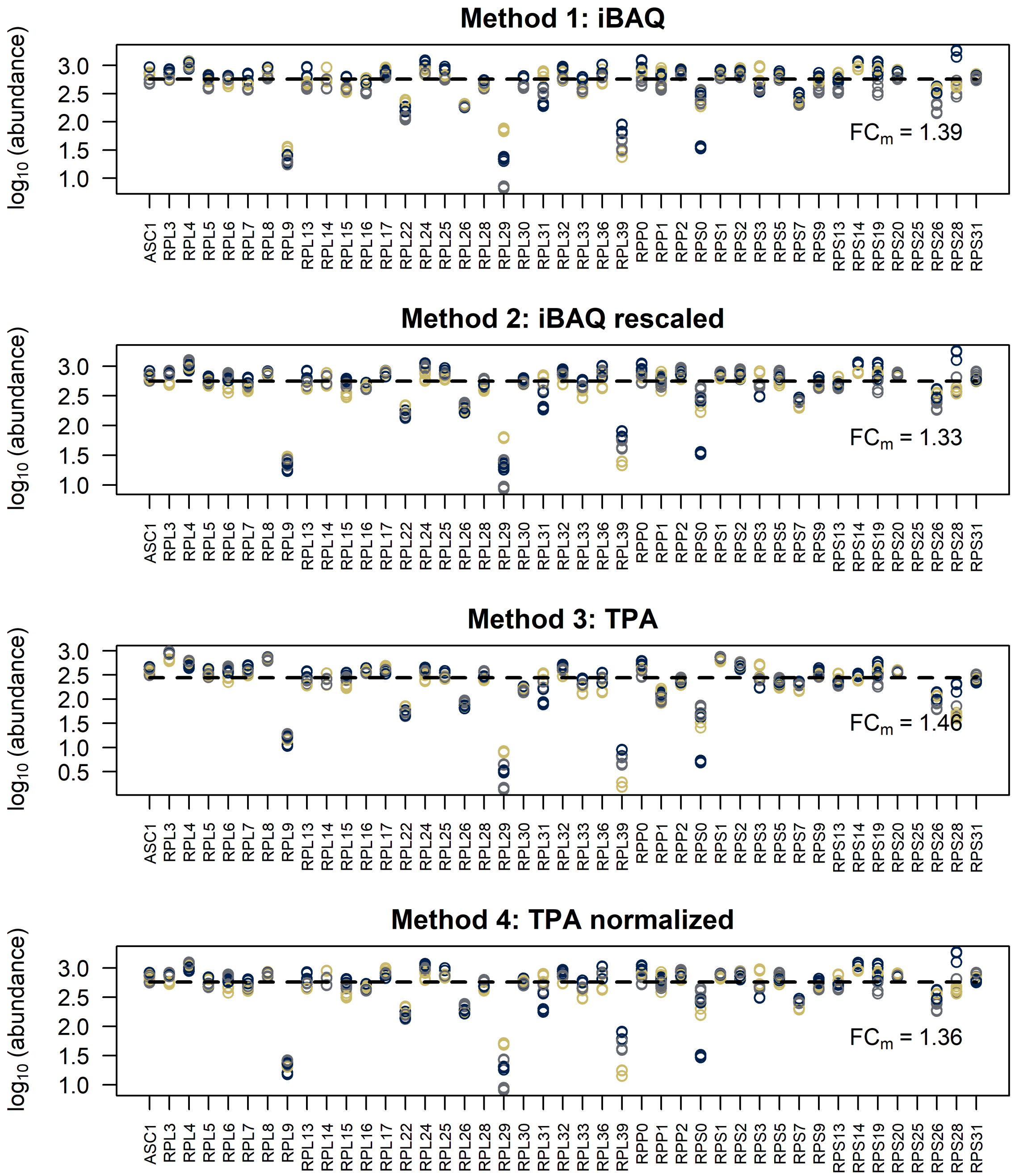

Figure S4: Comparison of predicted ribosomal subunit abundances [fmol/sample], according to all methods. Colors correspond to different technical replicates. The median value is displayed with a segmented line, and the median fold change to that line for all data is displayed.

These distributions are not very different between them (with exception of method 3), as we can see in the cumulative distributions:

```
## [1] "UPS2 abundance error - number of FC compared = 167"
## [1] "UPS2 abundance error - number of FC compared = 167"
## [1] "UPS2 abundance error - number of FC compared = 167"
```

```
## [1] "UPS2 abundance error - number of FC compared = 166"
## [1] "UPS2 abundance error of iBAQ = iBAQrescaled: p-val = 0.352413929013194"
## [1] "UPS2 abundance error of iBAQ = TPA: p-val = 1.85823305409727e-07"
## [1] "UPS2 abundance error of iBAQ = TPAnorm: p-val = 0.486339715062478"
## [1] "UPS2 abundance error of iBAQrescaled = TPA: p-val = 1.85823305409727e-07"
## [1] "UPS2 abundance error of iBAQrescaled = TPAnorm: p-val = 0.980544617660828"
## [1] "UPS2 abundance error of TPA = TPAnorm: p-val = 7.58421095259365e-07"

## [1] "Ribosomal stoichiometry error - number of FC compared = 731"
## [1] "Ribosomal stoichiometry error - number of FC compared = 731"
## [1] "Ribosomal stoichiometry error - number of FC compared = 731"
## [1] "Ribosomal stoichiometry error - number of FC compared = 731"
## [1] "Ribosomal stoichiometry error of iBAQ = iBAQrescaled: p-val = 0.00357985056692356"
## [1] "Ribosomal stoichiometry error of iBAQ = TPA: p-val = 9.95671238679385e-07"
## [1] "Ribosomal stoichiometry error of iBAQ = TPAnorm: p-val = 0.010405685995723"
## [1] "Ribosomal stoichiometry error of iBAQrescaled = TPA: p-val = 1.82208803600759e-10"
## [1] "Ribosomal stoichiometry error of iBAQrescaled = TPAnorm: p-val = 0.485424882465365"
## [1] "Ribosomal stoichiometry error of TPA = TPAnorm: p-val = 1.47318354182246e-08"
```

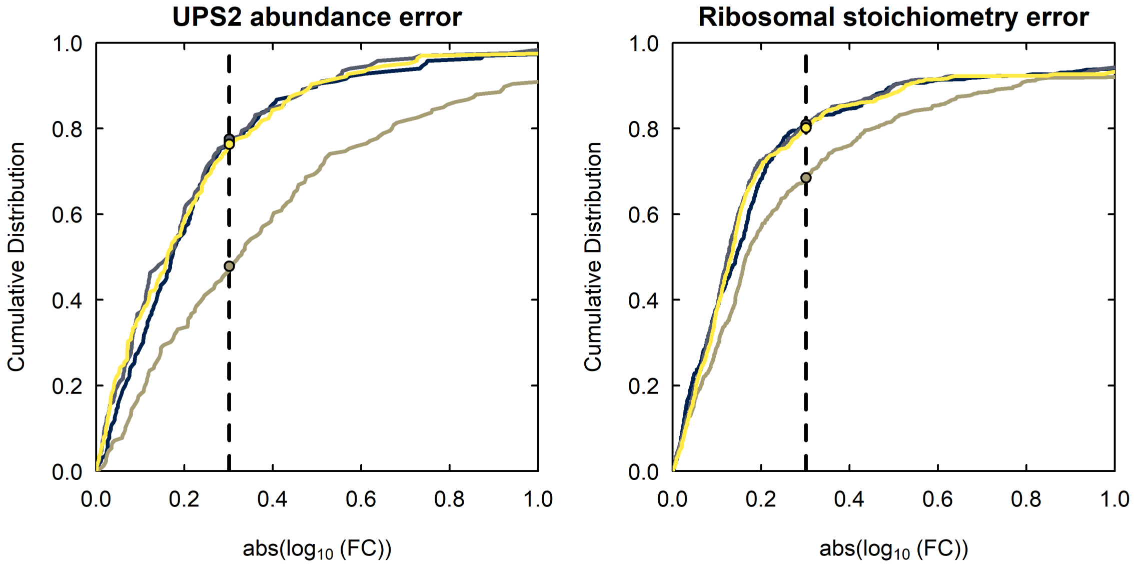

Figure S5: Cumulative distributions of absolute fold changes for both accuracy evaluation metrics: differences of predicted Vs experimental values of UPS2 (left) and differences to median value in ribosomal measurements (right). A fold change of 2 is indicated with a vertical segmented line. Colors represent the methods: 1) iBAQ (blue), 2) iBAQ rescaled (gray), 3) TPA (brown) and 4) TPA normalized (yellow).

##### 4.3 Evaluating precision

We now display the variability of the final abundance data between biological replicates and between batches, for all methods. For that we first define a function that gives all possible combinations between replicates. For instance, for biological replicates, the text in the variable's label regarding biological replicate (`.R1.1`, `.R2.1` and `.R3.1`) will be first removed, and then 2 columns will be paired up if the rest of the name matches (meaning it's the same batch/MS method but 2 different biological replicates). With that, we then plot the data with variability plots and a PCA:

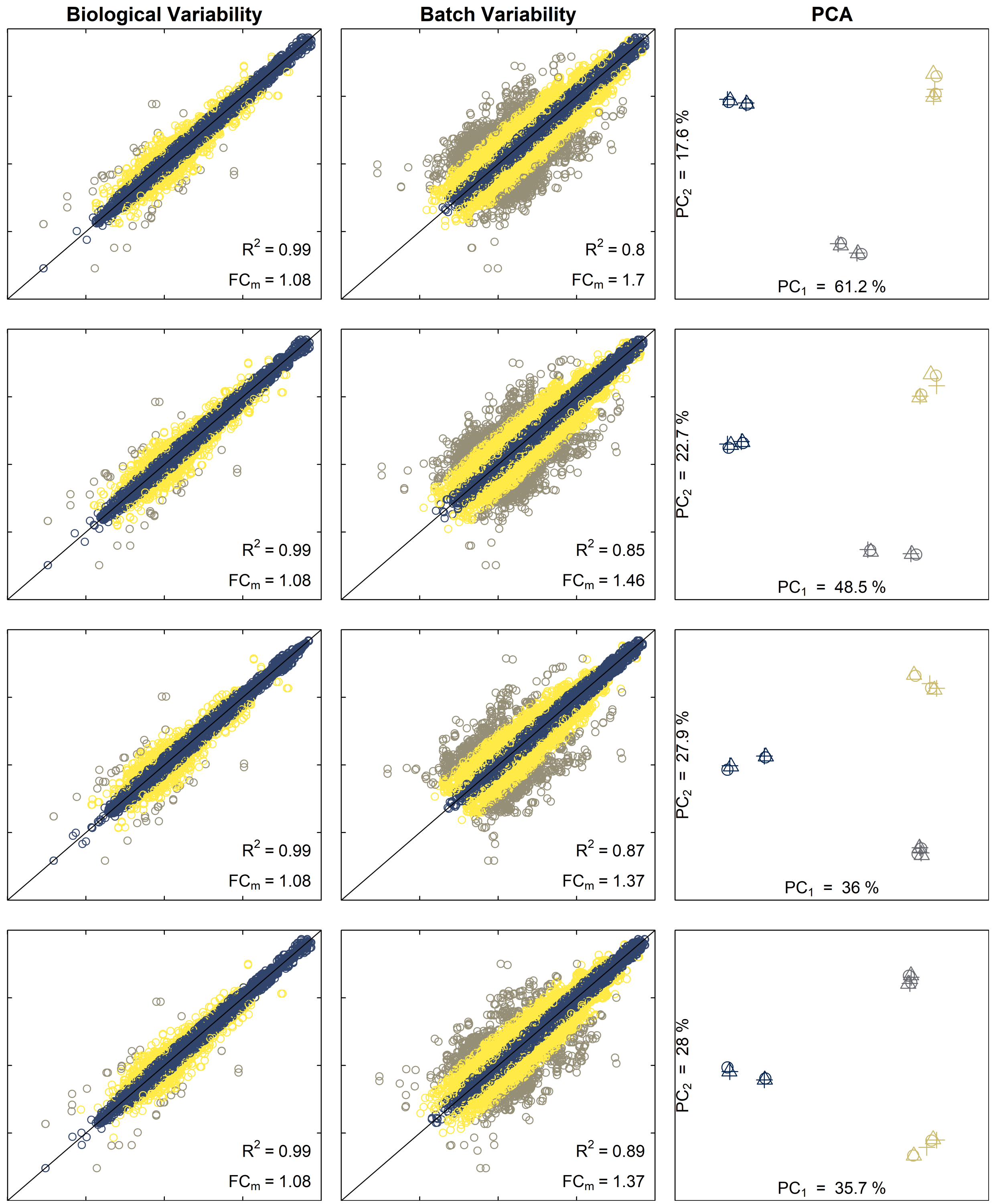

Figure S6: Comparison of data variability from 1) iBAQ (1st row), 2) rescaling iBAQ (2nd row), 3) TPA (3rd row) and 4) normalized TPA (4th row). In the variability plots (left and middle columns, log10(abundance [fmol/sample]) both in the x-axis and y-axis), 2 abundance values for a given protein are plotted if they belong to the same biological replicate or batch, respectively. Blue is a FC lower than 2, yellow between 2 and 10, and gray over 10. In the PCA plots (right column), colors refer to MS batches and shapes to biological replicates.

We see a lower median fold change between batches + a better separation of the "batch clusters" in the PCA when we use method 3 or 4 (PC1+PC2 represents less variability). This means that by using rescaled MS data (methods 3 or 4) we achieve lower variability between batches than with methods 1 and 2. This is confirmed by looking at the breakdown of proteins by method:

Table S1: Protein breakdown by method and type of replicate. For each protein, the maximum fold change is considered.

| | Method 1 | Method 2 | Method 3 | Method 4 |
|---|---|---|---|---|
| **Variability between biological replicates:** | | | | |
| FC $<$ 2 | 89.4% | 89.4% | 89.4% | 90.8% |
| 2 $<$ FC $<$ 10 | 9.7% | 9.7% | 9.7% | 8.4% |
| 10 $<$ FC | 0.9% | 0.9% | 0.9% | 0.7% |
| **Variability between batches:** | | | | |
| FC $<$ 2 | 35.3% | 46.3% | 51.4% | 53.7% |
| 2 $<$ FC $<$ 10 | 53.4% | 45.3% | 41.9% | 40.2% |
| 10 $<$ FC | 11.3% | 8.4% | 6.7% | 6% |
| **Variability between all replicates:** | | | | |
| FC $<$ 2 | 29.5% | 40.2% | 45.1% | 48.1% |
| 2 $<$ FC $<$ 10 | 56.8% | 49.2% | 46.4% | 44% |
| 10 $<$ FC | 13.7% | 10.7% | 8.5% | 7.8% |

##### 4.3.1 Evaluating inter-batch precision

Let's look further into the reduction of batch variability, by looking at the cumulative distribution of batch variability for each method:

```
## [1] "Inter-batch precision - number of FC compared = 21320"
## [1] "Inter-batch precision - number of FC compared = 21320"
## [1] "Inter-batch precision - number of FC compared = 21320"
## [1] "Inter-batch precision - number of FC compared = 21061"
## [1] "Inter-batch precision of iBAQ = iBAQrescaled: p-val = 0"
## [1] "Inter-batch precision of iBAQ = TPA: p-val = 0"
## [1] "Inter-batch precision of iBAQ = TPAnorm: p-val = 0"
## [1] "Inter-batch precision of iBAQrescaled = TPA: p-val = 0"
## [1] "Inter-batch precision of iBAQrescaled = TPAnorm: p-val = 0"
## [1] "Inter-batch precision of TPA = TPAnorm: p-val = 0.946855360017903"
```

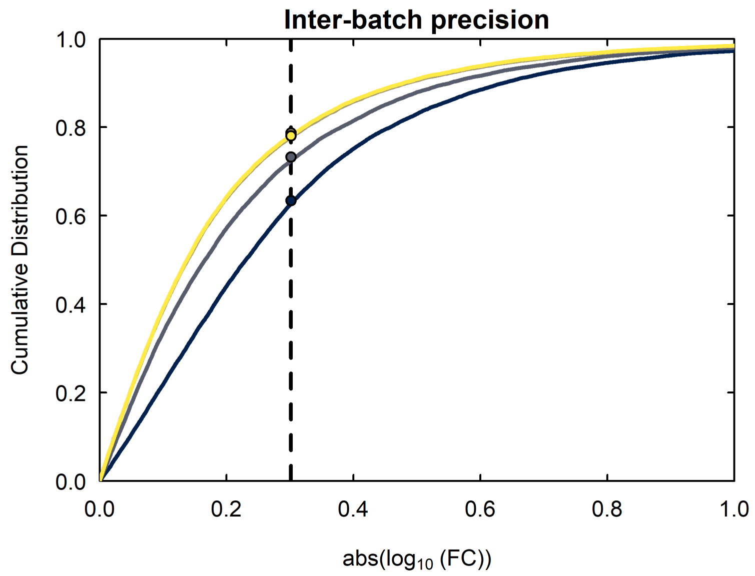

Figure S7: Fold change cumulative distributions of batch variability for all methods. A fold change of 2 is indicated with a vertical segmented line. Colors represent the methods: 1) iBAQ (blue), 2) iBAQ rescaled (gray), 3) TPA (brown) and 4) TPA normalized (yellow).

We can also look at batch variability by plotting each FC to the corresponding abundance, together with a "UPS2 window" that shows the abundance levels that are detected by the UPS2:

```
## [1] "Method 1: iBAQ -> 197 proteins below UPS2 detection range"

## [1] "Method 2: iBAQ rescaled -> 288 proteins below UPS2 detection range"

## [1] "Method 3: TPA -> 73 proteins below UPS2 detection range"

## [1] "Method 4: TPA normalized -> 234 proteins below UPS2 detection range"
```

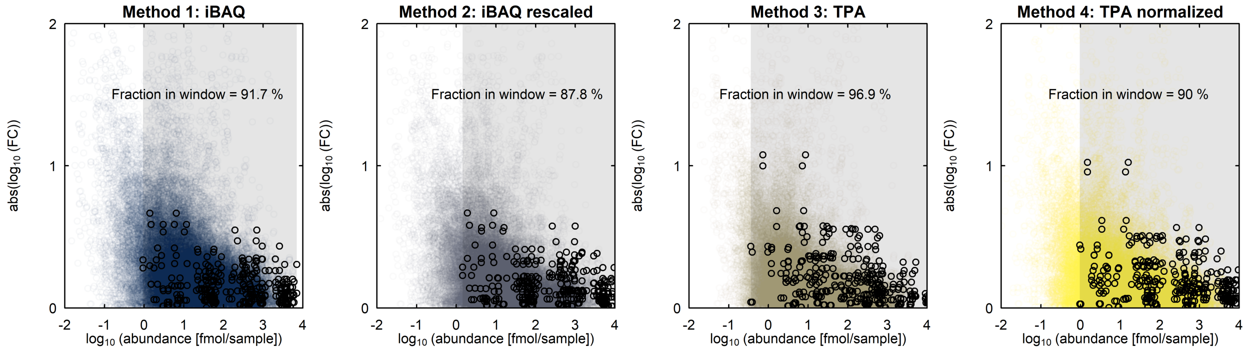

Figure S8: Fold change Vs abundances for all methods. The detection window of UPS2 and the UPS2 datapoints are highlighted in gray and black, respectively.

We see that in all cases more than 85% of the data falls within the detection range of UPS2, which is good. However, all datasets look similar in shape, so instead let's look at the trend of the data with the help of smooth splines:

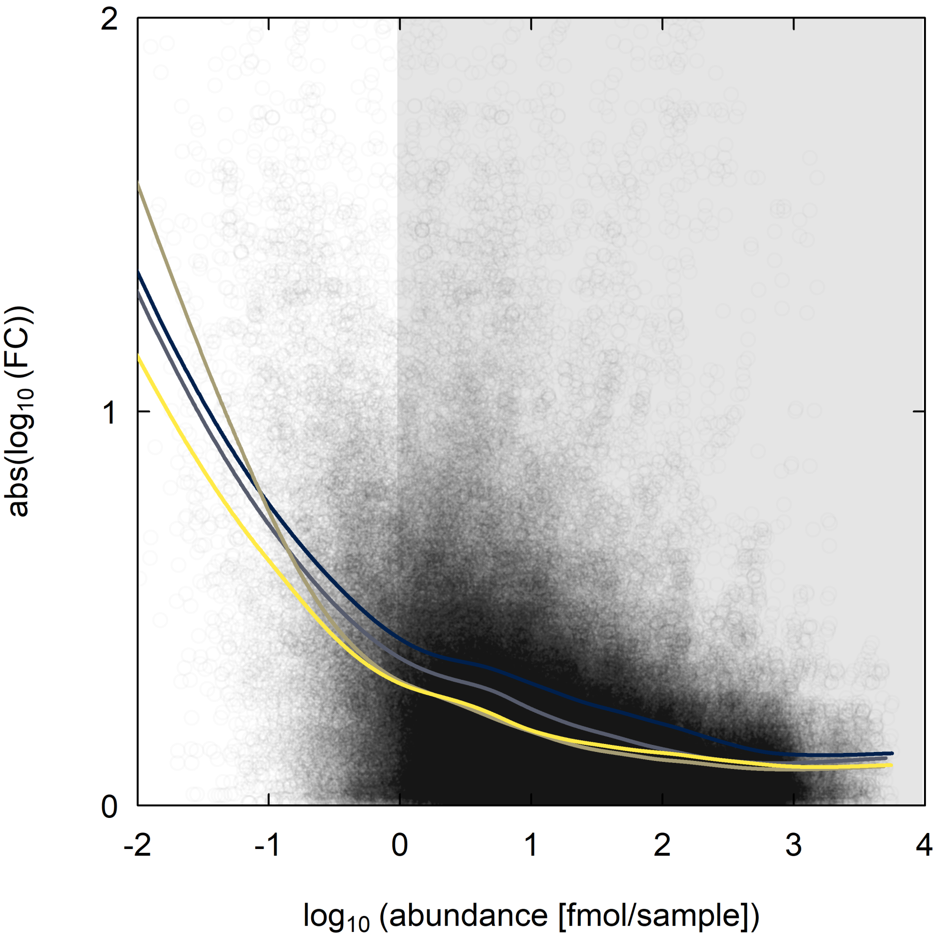

Figure S9: Fold change Vs abundances for all methods + smoothing splines. Colors represent the methods: 1) iBAQ (blue), 2) iBAQ rescaled (gray), 3) TPA (brown) and 4) TPA normalized (yellow). The detection window of UPS2 is highlighted in gray.

We see that method 4 (normalized TPA) has overall less variability than the both iBAQ and rescaled iBAQ, both for lowly and highly abundant proteins. It also performs better than the traditional TPA (method 3) at low abundances.

#### 4.4 Additional comparisons

##### 4.4.1 ES curves

Let's take a look at the ES real abundance values Vs the normalized intensity data from method 4, and compare the "linear model" mentioned in section 3.4 (method 4) to a linear fit to the data (method 1):

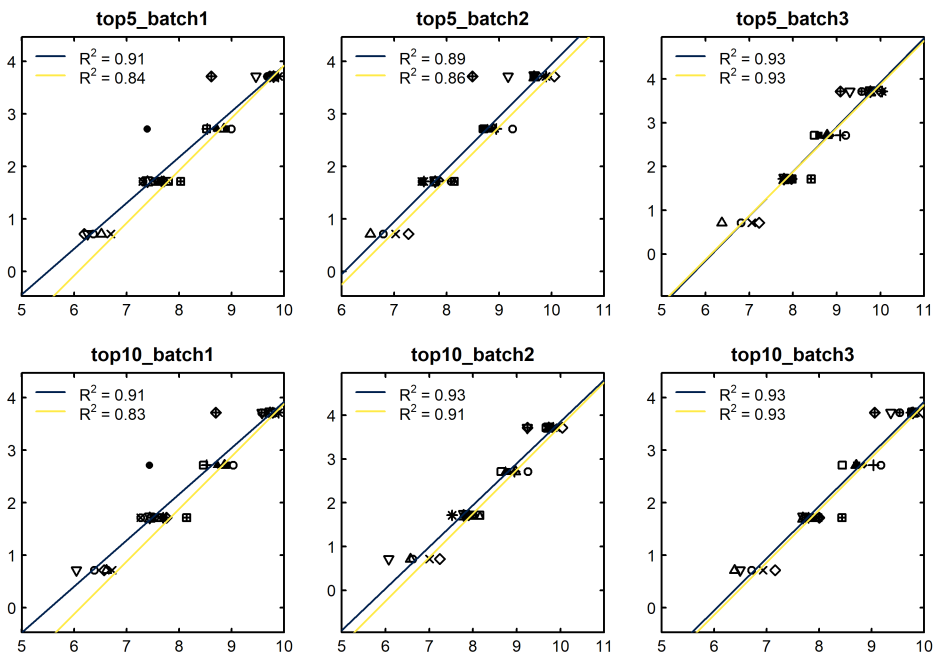

Figure S10: log10(abundance [fmol/sample]) Vs log10(normalized MS intensity) of the 30/48 UPS2 proteins that were detected and measured by the MS, together with 2 linear models used later for converting the data: method 1, iBAQ (blue); and method 4, rescaling the normalized intensities (yellow). Within each of the 4 orders of magnitude, each symbol corresponds to a different protein.

The blue fits give us the transformation from light (L) MS intensity of the UPS2 proteins to abundance (fmol/sample) with iBAQ. But they don't look the best (considering they are in log10 space), many other curves (as the yellow ones) can almost equally well fit that data. Let's see everything in the same plot:

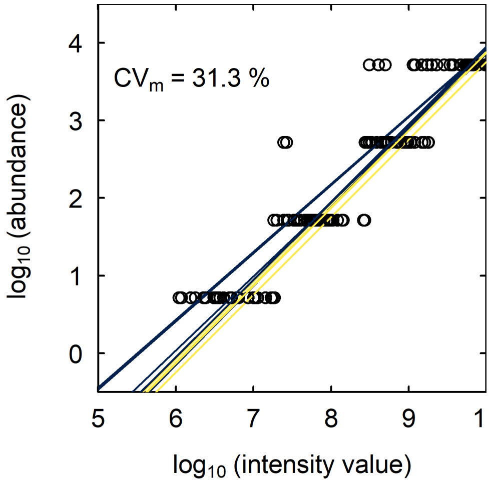

Figure S11: log10(abundance [fmol/sample]) Vs log10(normalized MS intensity) of the 30/48 UPS2 proteins that were detected and measured by the MS, together with 2 linear models used later for converting the data: method 1, iBAQ (blue); and method 4, rescaling the normalized intensities (yellow). Average coefficient of variation (CVm) within each protein is shown in the upper left corner.

In conclusion, we can skip entirely the UPS2 data and just assume that the normalized MS intensities should always adds up to a given protein amount. With this, we can reproduce very closely the ES curves, and achieve more consistent results across samples. In this approach, the iBAQ run is only used to do the rescaling, but this could be equally performed with any SILAC run.

###### 4.4.2 Predictions between methods

Plotting the data between methods shows us that predictions are not very different:

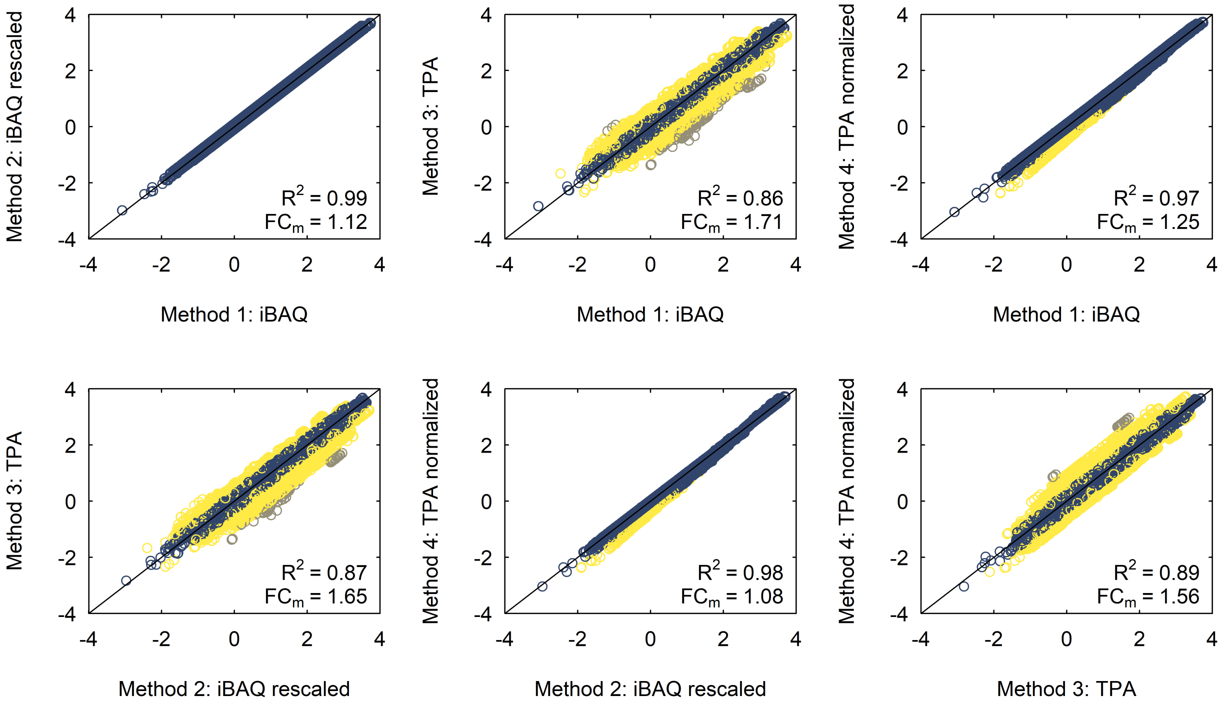

Figure S12: Comparison of predictions [fmol/sample] between all methods (on log10 scale). Blue is a FC lower than 2, yellow between 2 and 10, and gray over 10.

###### 4.4.3 Sequence length

Finally, let's see the predicted abundances of all samples compared to sequence length:

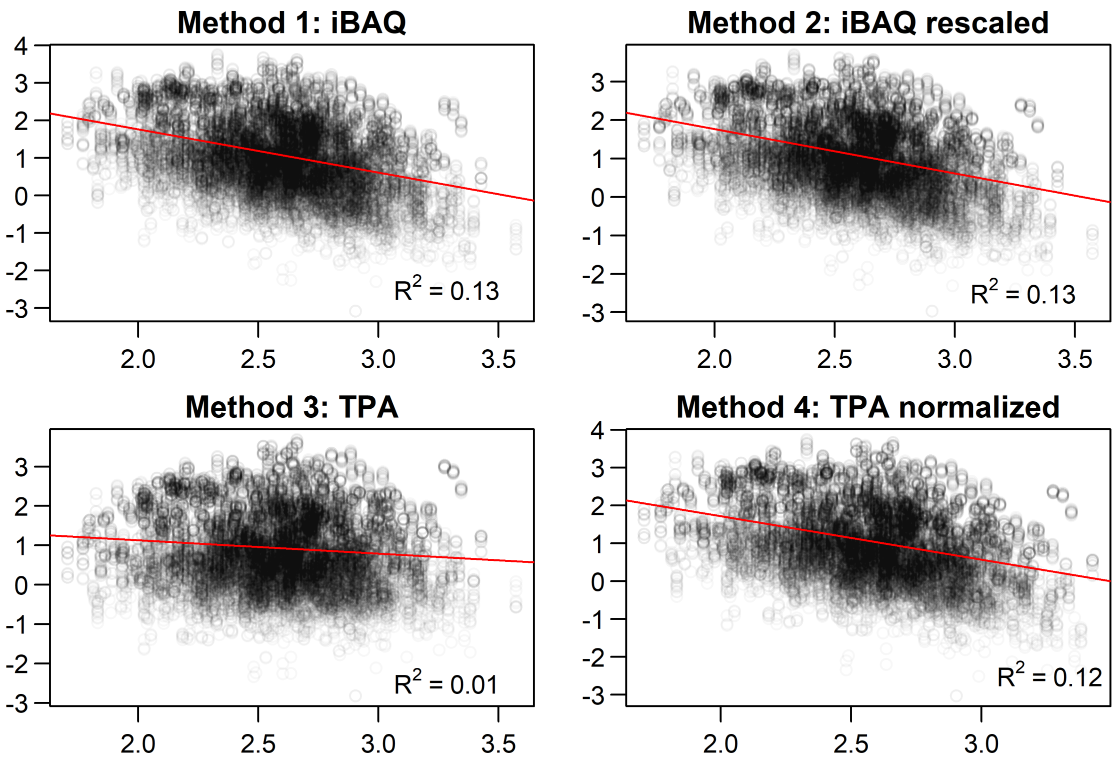

Figure S13: Predicted abundances [fmol/sample] Vs sequence length (both in the log10 space) for all proteins and all methods.

We can see that methods 1, 2 and 4 all have similar correlation values, while method 3 correlates less than the other 3 methods to protein length. This is expected, because methods 1, 2 and 4 all normalize by the number of theoretical peptides, which correlates well with sequence length:

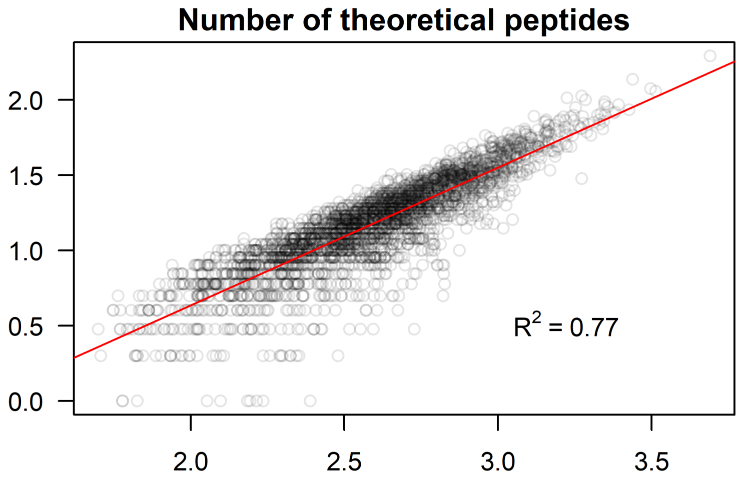

Figure S14: Number of theoretical peptides Vs sequence length for all proteins (both in the log10 space).

With this we also see that sequence length can work as a good proxy for the number of theoretical peptides (if the latter is not available).

### References

Cox, Jürgen, and Matthias Mann. 2008. "MaxQuant enables high peptide identification rates, individualized p.p.b.-range mass accuracies and proteome-wide protein quantification." *Nature Biotechnology* 26 (12): 1367–72. https://doi.org/10.1038/nbt.1511.

Jenner, Lasse, Sergey Melnikov, Nicolas Garreau de Loubresse, Adam Ben-Shem, Madina Iskakova, Alexandre Urzhumtsev, Arturas Meskauskas, Jonathan Dinman, Gulnara Yusupova, and Marat Yusupov. 2012. "Crystal structure of the 80S yeast ribosome." Elsevier Current Trends. https://doi.org/10.1016/j.sbi.2012.07.013.

Schwanhäusser, Björn, Dorothea Busse, Na Li, Gunnar Dittmar, Johannes Schuchhardt, Jana Wolf, Wei Chen, and Matthias Selbach. 2011. "Global quantification of mammalian gene expression control." *Nature* 473 (7347): 337–42. https://doi.org/10.1038/nature10098.

Wiśniewski, Jacek R., Paweł Ostasiewicz, Kamila Duś, Dorota F. Zielińska, Florian Gnad, and Matthias Mann. 2012. "Extensive quantitative remodeling of the proteome between normal colon tissue and adenocarcinoma." *Molecular Systems Biology* 8 (611). https://doi.org/10.1038/msb.2012.44.

Wiśniewski, Jacek R., and Dariusz Rakus. 2014. "Multi-enzyme digestion FASP and the 'Total Protein Approach'-based absolute quantification of the Escherichia coli proteome." *Journal of Proteomics* 109: 322–31. https://doi.org/10.1016/j.jprot.2014.07.012.
